# Characterization of the gut mycobiome in patients with non-alcoholic fatty liver disease and correlations with serum metabolome

**DOI:** 10.1101/2025.03.10.642414

**Authors:** Ning Zheng, Dangrang Wang, Guorui Xing, Yidan Gao, Shenghui Li, Jin Liu, Jian Kang, Shanshan Sha, Lin Cheng, Shao Fan, Jian Yu, Qiulong Yan, Chunmeng Jiang

## Abstract

**Background:** The relationship between the gut mycobiome and non-alcoholic fatty liver disease (NAFLD) is largely unexplored.

**Methods:** In this study, we used a publicly available shotgun metagenomic sequencing dataset of fecal samples to compare the gut fungal communities of 90 NAFLD patients with those of 90 healthy controls. We examined their correlations with gut bacterial communities, host serum metabolite s, and the interactions of key taxa within these networks.

**Results:** Our results revealed a remarked dysbiosis in the gut mycobiome of NAFLD patients, characterized by a significant increase in four fungal taxa, including *Pseudopithomyces sp*. c174, *Mucor sp*. c176, *Aspergillus sp*. c25, and *uncultured Ascochyta* c213. Multi-omics analysis showed that the gut mycobiome significantly contributes to the host serum metabolome. *Pseudopithomyces s p*. c174 was notably associated with some beneficial blood metabolites, such as glycoursodeoxycholic acid and alpha-linolenic acid, whereas the NAFLD-enriched pro-inflammatory phenylacetic acid was linked to *Aureobasidium sp*. c170 and *Basipetospora sp*. c193. Network analysis revealed significant alterations in the gut microbiome of NAFLD patients, highlighting the potential roles of *Alternaria alternata* c42, *Penicillium sp*. c5, and *uncultured Malassezia* c303. Moreover, predictive models that integrated both bacterial and fungal taxa improved prediction accuracy, emphasizing the crucial role of gut fungi in NAFLD.

**Importance:** Non-alcoholic fatty liver disease (NAFLD) is a widespread liver condition with limited treatments. While gut bacteria have been extensively studied in NAFLD, the role of gut fungi (the mycobiome) remains poorly understood. This study fills that gap by characterizing gut fungal communities in NAFLD patients and linking these changes to their blood metabolite profiles. We found that NAFLD patients show distinct fungal imbalances, which correlate with key metabolic alterations in the host. These findings expand our understanding of NAFLD by spotlighting gut fungi as important contributors to disease progression. This insight opens new avenues for potential biomarkers and therapies targeting the gut-liver axis.

**Conclusions:** Our findings provide a comprehensive understanding of the link between the gut mycobiome and NAFLD progression, which is essential for guiding future therapeutic research.

## Introduction

Non-alcoholic fatty liver disease (NAFLD) is the most common chronic liver disease worldwide, affecting about one-quarter of the global population [1]. The disease includes a spectrum from the relatively mild non-alcoholic fatty liver (NAFL) to the more severe non-alcoholic steatohepatitis (NASH), which can progress to cirrhosis and hepatocellular carcinoma (HCC) associated with NAFLD (NAFLD-HCC) [2]. This progression represents a significant public health issue due to the increased risk of morbidity and mortality. Although the rising prevalence of NAFLD is closely linked with the global obesity epidemic [3], it also occurs in non-obese individuals, indicating diverse underlying causes [4].

Contemporary studies indicate that the multifactorial causes of NAFLD, include genetic factors, obesity, an imbalanced gut microbiome, microbial metabolites, and weakened gut barrier function [5, 6]. Among, the increasing evidence points to the gut microbiome as a crucial element in NAFLD development and progression, linking diet, metabolism, and liver health [7].

The gut microbiome, a complex ecosystem with over 100 trillion microorganisms like bacteria, fungi, viruses, and archaea, plays a vital role in human health. Changes in its composition can worsen metabolic imbalances and inflammation, increasing the severity of various diseases. For instance, gut microbiome dysbiosis is linked to increased gut permeability, allowing bacterial products to reach the liver through the portal vein, like lipopolysaccharides, and secondary bile acids [8], thus promoting inflammation and fibrosis [4, 9]. Similarly, a decrease in beneficial bacterium (*Bifidobacterium*) and an overgrowth of harmful fungus (*Candida sp*.) have been observed in NAFLD patient s, with lower bacterial diversity seen in more severe cases [10]. Overall, the gut microbiome appears to significantly influence NAFLD health outcomes.

Due to bacteria being the most abundant species in the gut, much research has primarily focused on the bacterial component of the gut microbiome. However, recent studies emphasize the significant role of the gut mycobiome, a complex fungal network in the gastrointestinal tract, in NAFLD [11]. These studies show distinct fungal profiles in NAFLD patients, suggesting the gut mycobiome could be a new biomarker for diagnosing and monitoring disease progression [11]. For example, specific fungal taxa changes have been linked to the severity of liver disease in non-obese NAFLD patients [12], underlining the importance of understanding gut mycobiome dynamics in NAFLD. Antifungal treatments have shown promise in ameliorating NAFLD markers in experimental models, further supporting the potential therapeutic target within the gut mycobiome [12]. Additionally, certain fungal species, like C. albicans, significantly affect immune responses and metabolic pathways related to liver diseases. *C. albicans* secretes candidalysin, which stimulates IL-1α, IL-1β, I L-8, IL-36, and activates the NLRP3 inflammasome [13]. This activation is critical in the advancement of metabolic diseases including obesity, type 2 diabetes mellitus (T2DM), and NAFLD [14]. Thus, understanding the specific roles and interactions of gut fungi in NAFLD could uncover new mechanisms of disease progression and potential intervention targets.

This study aims to explore the role of the gut mycobiome in NAFLD and its correlations with serum metabolome. By analyzing gut mycobiome compositions in 90 healthy controls and 90 NAFLD patients through comprehensive multi-omics approaches [15], including fecal mycobiome profiling and serum metabolomics, this research seeks to identify specific fungal markers associated with NAFLD. Additionally, investigating the interactions between gut fungi and host metabolism may provide mechanistic insights into the mycobiome’s contribution to NAFLD. This integrative study aims not only to enhance our understanding of the gut mycobiome’s role in NAFLD but also to explore new diagnostic and therapeutic strategies targeting the complex interplay between the gut microbiome, liver health, and metabolism.

## Methods

### Data source and processing

All fecal metagenomic sequencing samples from 90 NAFLD patients and 90 healthy individuals used in this study were downloaded from the NCBI Sequence Read Archive (SRA) under the accession ID PRJNA728908 and PRJNA686835 (only 4 control samples with ID SRR13279648, SRR1 3279753, SRR13279664, and SRR13279666) [15]. To ensure data quality, we employed fastp [16] to process each metagenomic sample. The raw metagenomic reads suffered from several filtering steps, including trimming of polyG tails and removal of low-quality reads as follows: (1) reads shorter than 90bp; (2) reads with a mean Phred quality score lower than 20; (3) reads with over 30% of their bases having a Phred quality score lower than 20; (4) reads with a mean complexity below 30%; and (5) unpaired-end reads. Additionally, the serum metabolomic profiles of all subjects were downloaded from the Metabolite database under the accession ID MTBLS2615.

### Taxonomic profiling of the gut mycobiome

Based on the methodologies outlined in our previously published articles [17, 18], we have updated the original database by incorporating a total of 229 new genomes from the National Center of Biotechnology Information (NCBI) database. These strains were then clustered into 302 non-redundant human-associated fungi species using an average nucleotide identity (ANI) threshold of 95%. To minimize the impact of non-specific mapping of reads to fungal genomes in subsequent analysis, we mapped the quality-faltered reads against three databases: the GRCh38 genome, the Unified Human Gastrointestinal Genome (UHGG) collection [19], and the SILVA rRNA database [20]. This step allowed us to exclude reads derived from human or prokaryotic sources. For each sample, the remaining reads were aligned against our customized catalog of gut fungal genomes using bow tie2 [21], and the read counts for each genome were calculated. To generate mycobiome composite on profiles, the read count of each genome was first normalized by dividing its genomic size, and the normalized read count was further divided by the sum of all normalized read counts in a sample. This process defends the relative abundance of each population in the sample. For different fungal taxa, the relative abundance of a taxon was calculated as the sum of the relative abundance of all populations assigned into that taxon.

### Statistical analyses

Statistical analysis and visualization were carried out by the R platform. A Bray-Curtis distance matrix was generated using the square-root transformed the species-level taxonomic profiles, using the *vegdist* function from the *vegan* package [22]. Principal coordinates analysis (PCoA) was then performed on the distance matrix using the *pcoa* function in the *ape* package. We calculated the number of observed species by counting the species with a relative abundance greater than zero in e ach sample. Shannon’s and Simpson’s diversity indices were defined using the function *diversity* in the *vegan* package. The Wilcoxon rank-sum test and Student’s t-test were implemented using the f unction *wilcox*.*test* and *t*.*test* functions, respectively.

Permutational multivariate analysis of variance (PERMANOVA) was conducted using the *adonis* function in the *vegan* package, based on the Bray-Curtis distance matrix. The evaluation of explanatory power between different omics datasets was performed using stepwise PERMANOVA analysis, according to existing methods. For example, in the assessment of the explanatory power of the gut mycobiome on the serum metabolome, the following steps were followed: (1) The R-squared value (R^2^) for each fungal species with respect to the serum metabolome was calculated using the *adonis* function, and the resulting R^2^ was adjusted using the *RsquareAdj* function. (2) The fungal species exhibiting the largest adjusted R^2^ was selected as the first variate. (3) A second PERMANOVA analysis was executed using the first variate and each remaining fungal species as variables. (4) If the largest adjusted R^2^ from the second PERMANOVA analysis is smaller than that from the first PERMANOVA analysis, the latter is considered as the explanatory power of the gut mycobiome on the serum metabolome. (5) If the largest adjusted R^2^ from the second PERMANOVA analysis i s greater, the process is repeated for a third PERMANOVA analysis. In this case, the analysis includes another fungal species in addition to the two variates with the largest adjusted R^2^ from the second PERMANOVA analysis. (6) The process continues until the largest adjusted R^2^ from the last PERMANOVA analysis is smaller than that from the previous PERMANOVA analysis. The latter is then considered as the explanatory power of the gut mycobiome on the serum metabolome.

We performed a correlation analysis between the relative abundance of gut fungal species and the level of host metabolites using the function *cor*.*test* with the option ‘method=spearman’. The resulting p-values were then adjusted using the function *p*.*adjust* with the option ‘method=BH’. A correlation was considered significant if the adjusted p-value was less than 0.05.

The random forest classifier based on the gut mycobiome was built using the *randomForest* function followed by 5 times of five-fold cross-validations, and their performances were evaluated based on the area under the receiver operator characteristic curve (AUC) that was calculated by the ‘*roc*’ function. The importance ordering of markers was obtained via the ‘importance’ function.

The sunburst diagram of taxonomic hierarchy was generated using the function *plot_ly* in the pack age *plotly*. All other data were visualized using the function *ggplot* in the package *ggplot2*. The newly established iDIRECT approach was used to construct a bacterial co-occurrence network for each treatment, which efficiently eliminated indirect correlations and quantitatively inferred direct dependencies in a network [23]. To minimize false positives, only species detected in more than 10% biological replicates for each group were included. We focused on the interactions among taxa involved in bacteria and fungi.

## Results

### Sample information and fungal database

In this study, the composition of the gut mycobiome in NAFLD patients was investigated through a detailed re-analysis of deep-sequencing metagenomic samples that are publicly available. The dataset included samples from 90 NAFLD patients and 90 healthy controls (Table S1). In addition, we obtained serum metabolome profiles for 180 of the same subjects, allowing us to explore the potential association between the gut mycobiome and the development of NAFLD. To accurately define the mycobiome’s composition, we used a tailored fungal database utilizing strict filtering criteria (for detailed information, please consult the Methods section). This refined database includes 302 distinct reference fungal species, categorized based on a 95% ANI criterion, derived from an assortment of fungal genomes isolated from the human gut (Fig. 1). Subsequently, high-quality reads from each sample were aligned to the genomes of these 302 species present in our database, facilitating the creation of in-depth gut mycobiome profiles. Furthermore, our examination revealed the presence of 80 high-level taxa across the samples, which comprised 49 genera, 31 families, 19 orders, 11 classes, and 3 phyla (Fig. 1).

**Fig. 1.**
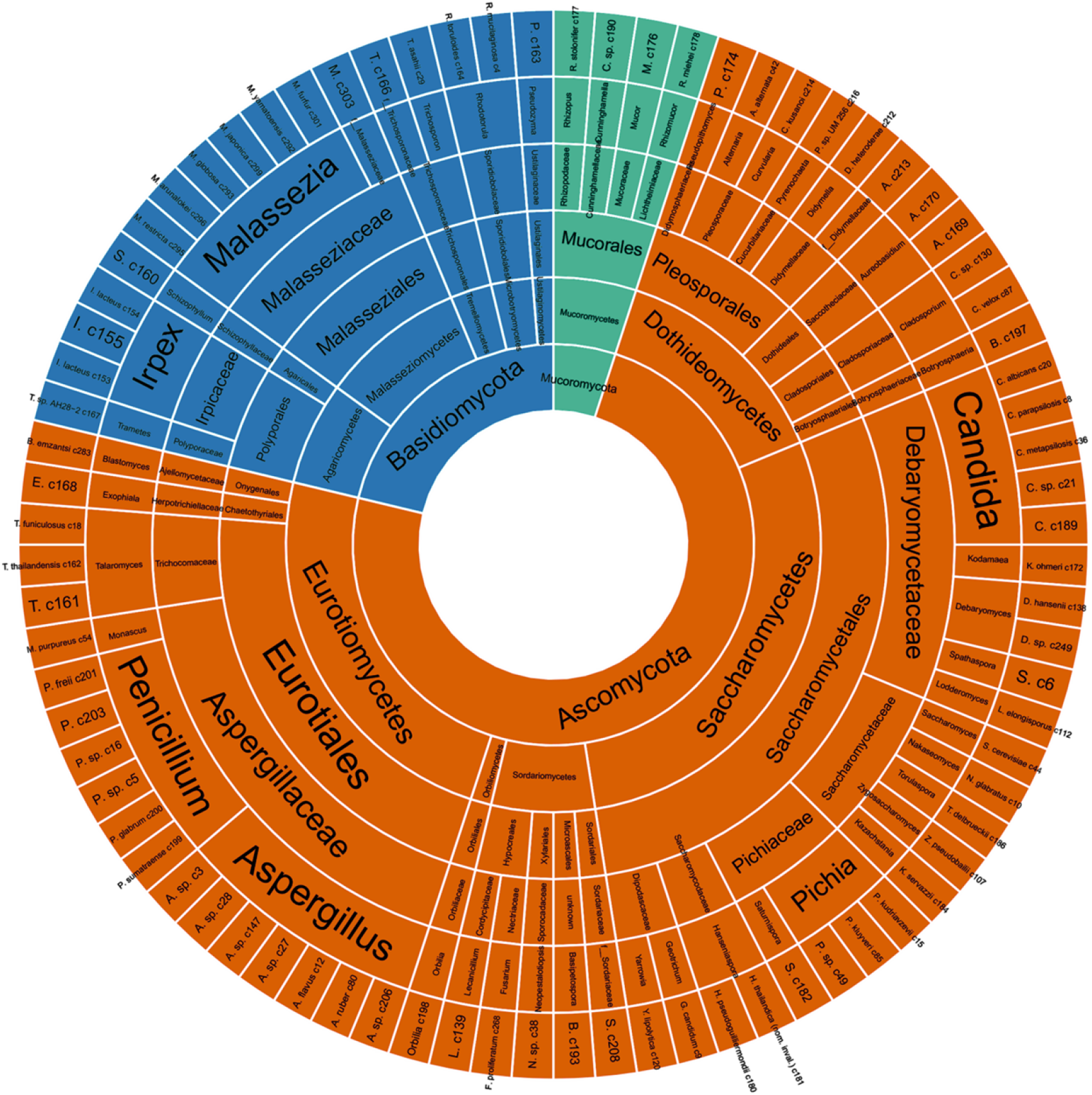
Sunburst diagram of taxonomic hierarchy for 302 gut fungal species and 80 high-level taxa.

### Altered gut mycobiome structure in NAFLD patients

We first compared the overall composition structure of the gut mycobiome between healthy controls and NAFLD patients using PCoA and PERMANOVA. PCoA based on Bray-cutis distance of species-level composition showed that the top two principal coordinate axes (PCoA1 and PCoA2) accounted for 29.9% and 25.1% of the total variation, respectively (Fig. 2a). Along the PCoA1, NAFLD patients showed a mild but statistically significant separation from healthy controls (Wilcoxon rank-sum test p = 0.14). PERMANOVA also showed a significant difference in the gut mycobiome between healthy controls and NAFLD patients (*adonis* p = 0.30).

**Fig. 2.**
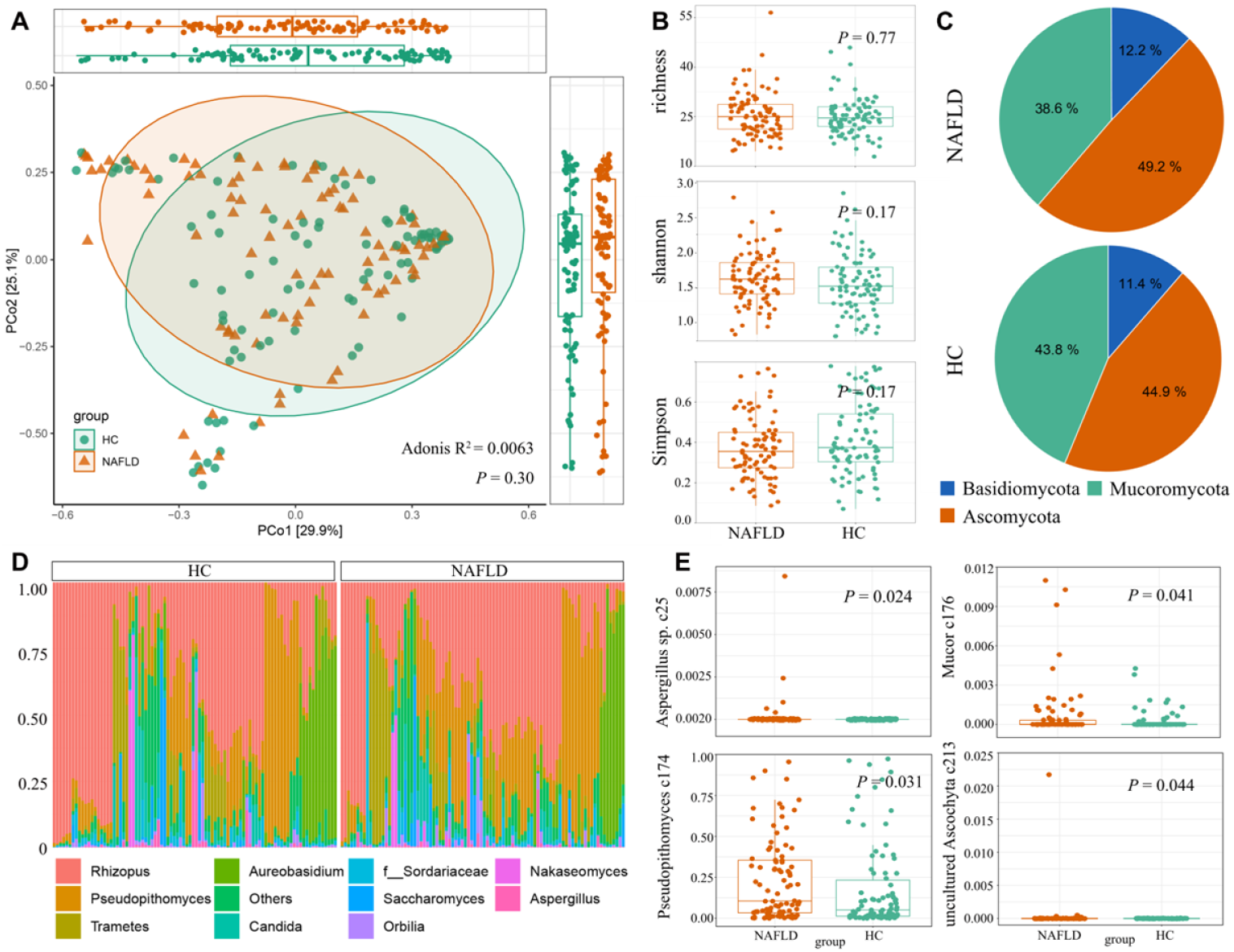
Comparison of gut mycobiome diversity and structure between NAFLD patients and healthy controls. **(A)** PCoA based on Bray–Curtis distance of the fungal profiles at the species level. The plot displayed the distribution of samples along PCoA1 and PCoA2, with ellipsoids indicating the 95% confidence interval for each group. The upper and right boxplots displayed the sample scores in PCoA1 and PCoA2. **(B)** Comparison of alpha diversity indexes between NAFLD patients and healthy controls. The p-value was determined by the Wilcoxon rank-sum test. **(C)** Pie chart showing the composition of fungal phyla in each group. The percentages represented the average relative abundance of each phylum. **(D)** Distribution of the top 10 abundant genera across all samples. **(E)** Boxplots showing the relative abundances of the species in each group. Statistical significance was determined using the Wilcoxon rank-sum test with Benjamini-Hochberg adjustment.

Alpha diversity metrics were employed to assess the gut mycobiome in NAFLD patients, utilizing three indices: richness, Shannon, and Simpson. However, the results indicated that there were no significant differences in the three indices of fungal alpha diversity between healthy controls and NAFLD patients (Wilcoxon rank-sum test, all p > 0.05; Fig. 2b).

In terms of the fungal taxa, the gut mycobiome of all subjects was usually dominated by Ascomycota, followed by Basidiomycota and Mucoromycota (Fig. 2c). At the genus level, *Rhizopus* was the most abundant species, while other common species, such as *Candida* and *Saccharomyces*, had relatively high abundances in both groups (Fig. 2d). Comparison analysis showed that 4 species were significantly enriched in NAFLD patients (Fig. 2e), including *Aspergillus sp*. c25 (Wilcoxon rank-sum test, adjusted p = 0.024), *Pseudopithomyces sp*. c174 (adjusted p = 0.031), *Mucor sp*. c176 (adjusted p = 0.041) and *uncultured Ascochyta* c213 (adjusted p = 0.044).

### Correlations between gut mycobiome and serum metabolome

In our study, we conducted an integrated analysis of multi-omics datasets, including the gut mycobiome and serum metabolome, to explore the potential contribution of the gut mycobiome to host health. The PERMANOVA analysis indicated that the gut mycobiome explained 38.2% of the variances in the host serum metabolome (Fig. 3a). When differentiating between healthy controls and NAFLD patients, both groups show considerable explanatory power of the gut mycobiome for the serum metabolic profiles. Specifically, the gut mycobiome of NAFLD patients and healthy control s accounted for 56.5% and 53.2% of the variance in the serum metabolome, respectively (Fig. 3b). To identify the fungal species associated with serum metabolites, we performed a correlation analysis using Spearman correlation analysis with Benjamini–Hochberg adjustment (q < 0.05). The results revealed that 24 fungal species were significantly correlated with at least one metabolite, including 1 NAFLD-enriched species (*Pseudopithomyces sp*. c174) (Fig. 3c). Among them, the NAFL D-enriched species *Pseudopithomyces sp*. c174 displayed significant positive correlations with 8 serum metabolites, namely 7-dehydrocholic acid, glycoursodeoxycholic acid, muro-cholic acid, alpha-linolenic acid, linoleic acid, heptadecanoic acid, nonadecanoic acid, and 3-hydroxybutyric acid. Through further comparative analysis, we identified 13 serum metabolites that exhibit significant differences between NAFLD patients and healthy controls. Among these, eight metabolites showed a clear correlation with fungal species, including beta-alanine, phenylacetic acid, isovaleric acid, valeric acid, methylglutaric acid, hydrocinnamic acid, and picolinic acid.

**Fig. 3.**
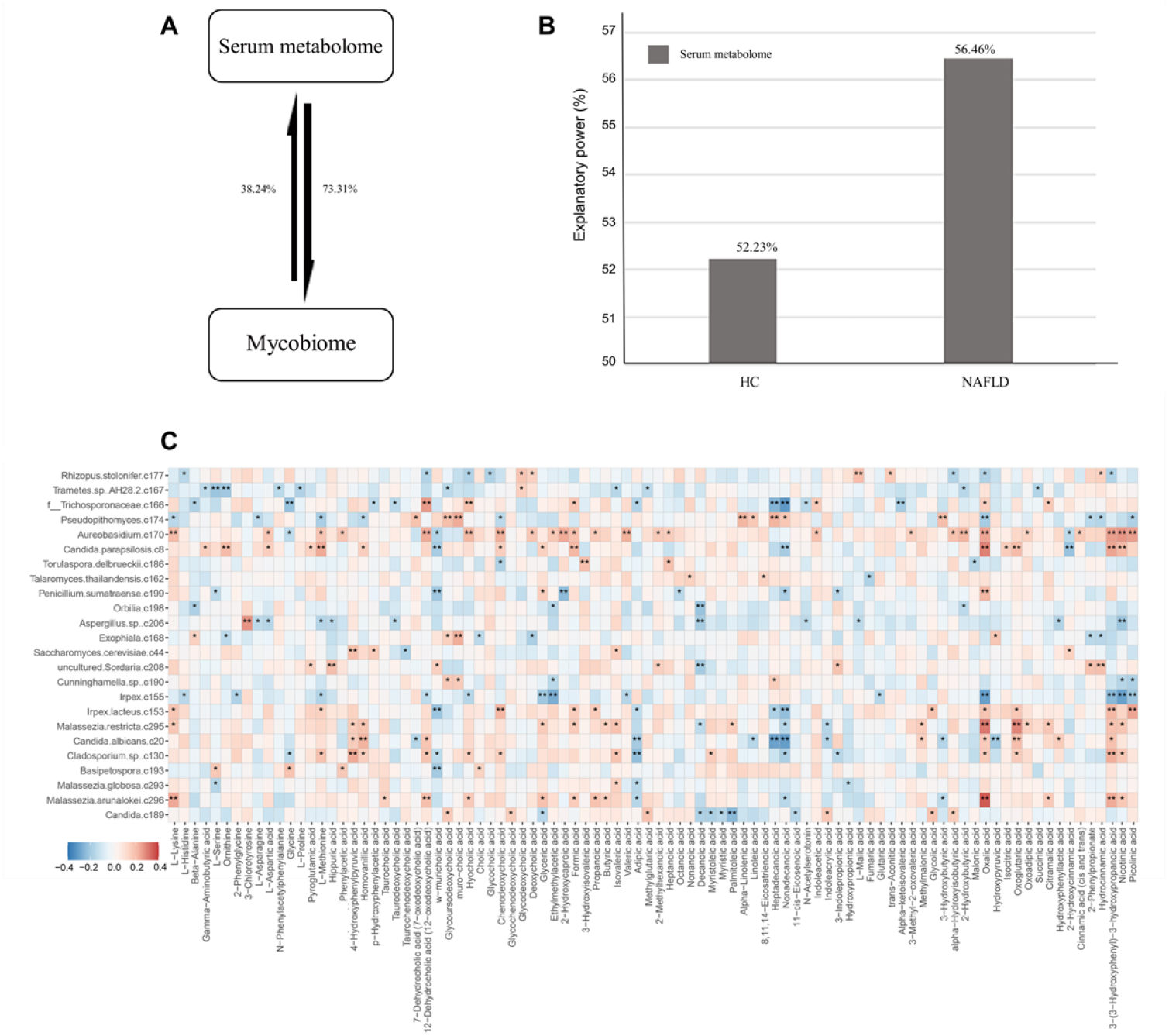
Interaction between the gut mycobiome and host serum metabolomes. **(A)** Explanatory power between different omics datasets. The explanatory power was quantified as adjusted R^2^ obtained from stepwise PERMANOVA analysis, as detailed in the Methods. Arrows indicated the direction and magnitude of explanatory power, with numbers indicating the corresponding values. **(B)** The explanatory power of the gut mycobiome on host serum metabolomes in NAFLD patients and healthy controls. **(C)** Heatmap displaying the Spearman’s rank correlation coefficients between fungal species and host metabolites. The q-value was obtained using the Benjamini and Hochberg ad justment. Correlations with a q-value<0.05 were considered statistically significant

### Comparative analysis of gut microbiome co-occurrence networks

Gut microbiome metabolism and associated biological activities are governed by a microbial consortium. Therefore, we employed co-abundance networks to link the taxa involved in this ecologically vital process, including both bacteria and fungi. We constructed co-abundance networks for the gut microbiome of both NAFLD patients and healthy control groups, ensuring that the correlation between any two nodes in the network exceeded 0.6. Compared to the gut microbiome network of the healthy control group, the network of NAFLD patients is more complex (Fig. 4a-b). Specifically, the gut microbiome network of NAFLD patients has significantly higher total nodes, total links, and average degree, while the average path distance is shorter. To further investigate the differences between the networks of NAFLD patients and healthy controls, we extracted key microbial taxa with connectivity greater than 10. In the gut microbiome network of the healthy controls, 31 species were identified, and in the network of NAFLD patients, another set of 33 species was identified (Fig. 4c-d). These key taxa connect more than 90% of the species within their respective networks. By comparing these key species, we found significant differences in fungi between the two gut microbiome networks. Specifically, in the gut microbiome network of NAFLD patients, eight fungal species played crucial roles, including *Curvularia kusanoi* c214, *Rhodotorula toruloides* c164, *Pseudozyma sp*. c163, *Alternaria alternata* c42, *Penicillium sp*. c5, *Blastomyces emzantsi* c283, *uncultured Malassezia* c303, and *Neopestalotiopsis sp*. c38. However, in the gut microbiome network of the healthy control group, *Schizophyllum sp*. c160, *Exophiala sp*. c168, *Aspergillus sp*. c27, *Rhodo torula toruloides* c164, and *Candida metapsilosis* c36 served as core elements.

**Fig. 4.**
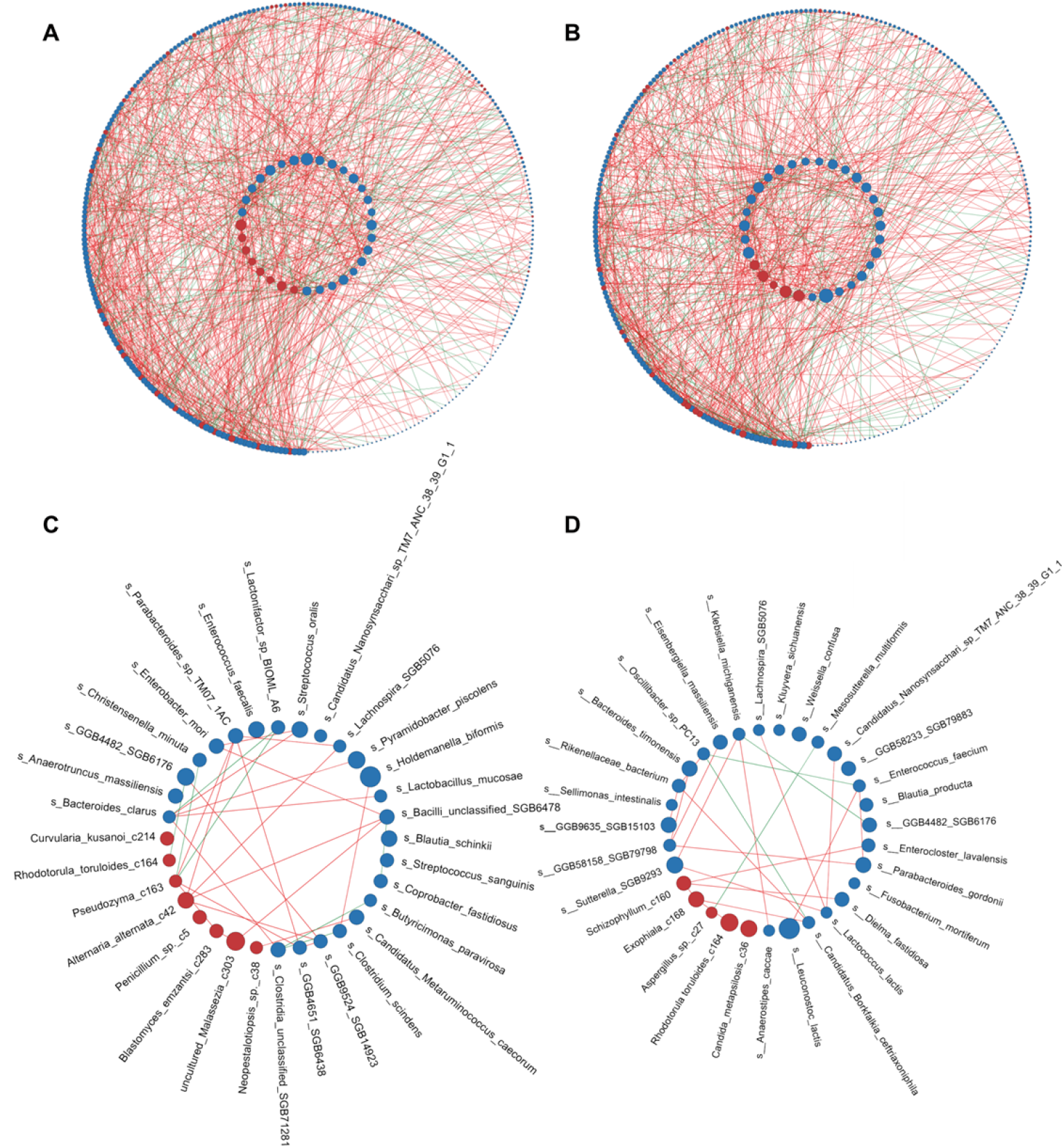
Correlation analysis among gut bacteriome and mycobiome. The network showed correlations between groups of gut bacteria and fungi in NAFLD patients **(A)** and HC group **(B)**, and extracted species with more than 10 connections for analysis **(C and D)**.

### Classification of NAFLD state based on the gut mycobiome

Finally, to evaluate the ability of the gut bacteria and mycobiome to classify NAFLD patients and healthy controls, we constructed a random forest model based on the relative abundances of the gut bacterial and fungal profiles. We found that both bacterial and fungal models achieved a cross-validated area under the curve (AUC) of 0.751 (95% confidence interval [CI] 0.719-0.782) and 0.618 (95% confidence interval [CI] 0.581-0.655), respectively (Fig. 5a). Compared to fungi, bacterial predictions were more pronounced in distinguishing between NAFLD patients and control groups. Interestingly, when we combined bacterial and fungal data for a joint prediction, we achieved a more significant result, with a cross-validated AUC of 0.772 (95% confidence interval [CI] 0.742-0.8 02), and both sensitivity and specificity were significantly higher than those achieved with single-taxa predictions. Although the majority of species in the prediction models are bacterial, these two fungi (*Alternaria alternata* c42 and *Candida parapsilosis* c8) played a significant role in further enhancing the models (Fig. 5b).

**Fig. 5.**
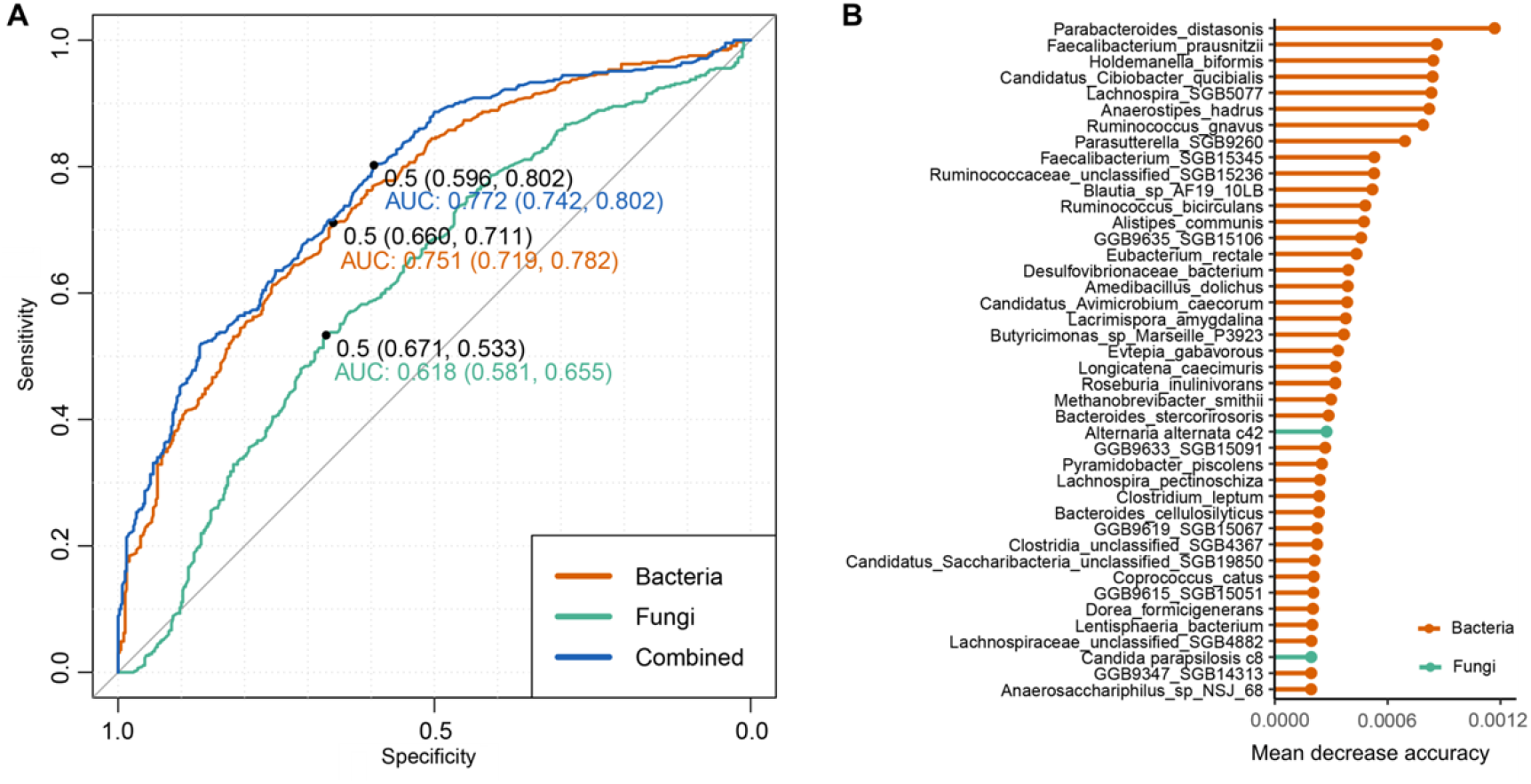
Classification of NAFLD status by the abundances of gut mycobiome and bacteriome. **(A)** ROC analysis for classification of NAFLD status using the gut bacterial and fungal signatures. **(B)** The 30 most discriminant signatures in the model classifying NAFLD patients and healthy controls. The bar lengths indicate the importance of the variable.

## Discussion

Numerous studies have shed light on the presence of gut dysbiosis in NAFLD patients, focusing primarily on analyzing the gut bacteriome [24, 25]. However, limited research has been conducted to investigate the relationship between the gut mycobiome and NAFLD [26]. Here, we used a comprehensive gut fungal genome database that closely relates to the human mycobiome. We explored the alterations in gut fungal communities in a cohort comprising 90 healthy controls and 90 NAFLD patients. Meanwhile, we investigated the relationships between gut fungal species and host serum metabolites, highlighting the potential impact of gut fungal species on NAFLD patients. Finally, based on the community characteristics of the gut microbiome, disease prediction was performed f or NAFLD patients, establishing potential biological markers.

We utilized a non-redundant gut fungal database containing 302 species-level genomes. Compared to the MetaPhlAn 4 database [27], which included 85 fungal species, our database has increased by 217 species, allowing for more accurate identification of fungal species within it. This database includes fungi known to colonize or infect various parts of the human body, while excluding fungi from dietary sources such as mushrooms and Ganoderma, as these fungi are typically transient and inactive in the gut. Utilizing this database, we observed that *Rhizopus* was the most prevalent genus in this cohort. Research shows that *Rhizopus* is one of the most obvious associations with NASH and is significantly linked to human diseases [28].

Microbiome analysis results show that there are no significant differences in fungal community diversity and structure between patients with NAFLD and healthy controls. We observed a slight increase in the alpha diversity index in NAFLD patients, consistent with a previous study on NAFLD based on the internal transcribed spacer 2 (ITS2) sequencing method [28]. Even though the results were not significant, they provide valuable insights into the trends that merit further investigation. Comparison analysis of fungal community composition revealed 4 species with a significant difference between NAFLD patients and healthy controls, with all four taxa enriched in the gut of the patients, including *Pseudopithomyces* c174, *Aspergillus sp*. c25, *Mucor* c176 and *uncultured Ascochyta* c213. These species have all shown significant changes in previous studies related to NAFLD. *Mucor sp*., opportunistic pathogens causing mucormycosis, are prevalent in patients with hematological cancers, uncontrolled diabetes, and are notably enriched in those with NAFLD [29]. However, previous studies indicated a significant reduction of *Pseudopithomyces* in NAFLD patients, but our results show the opposite trend. This discrepancy may be due to the expansion of the database, which allowed for the identification of new species, leading to new findings. Similarly, *Saccharomyces cerevisiae* c44, which is currently considered a potential probiotic in gut, did not show significant changes in this study. Given these new findings, future studies with larger sample sizes and more robust experimental designs are needed to clarify these findings.

In the past decade, research has highlighted the gut microbiome’s pivotal role in regulating host energy balance and metabolism [30]. Disruptions in its composition and function can impact metabol ism across various tissues, including the brain, adipose tissue, muscle, and liver. Compounds like lipopolysaccharides, secondary bile acids, and those resulting from carbohydrate and protein fermentation are closely linked to the gut microbiota-host metabolic axis and can contribute to metabolic disorders [31, 32]. Despite this strong association, the connection between fungal communities and host metabolites remains largely unexplored. Previous research has shown that although gut bacteria constitute over 80% of microbial sequences, their explanatory power on metabolic profiles is only 10% to 20% higher compared to the gut mycobiome [19, 33]. This indicates a significant impact of the mycobiome on metabolic profiles. Specifically, glycoursodeoxycholic acid is significantly correlated with NAFLD-enriched fungiand has been proven to effectively improve the condition [34]. It can alleviate endoplasmic reticulum stress in liver cells, a common issue in NAFLD, thereby reducing cell apoptosis and stabilizing calcium homeostasis. These effects contribute to helping prevent the progression of hepatic steatosis and inflammation, ultimately improving liver function in NAFLD patients. Similarly, alpha-linolenic acid provides antioxidant effects that reduce oxidative stress in liver tissue of NAFLD patients, offering hepatoprotective benefits and regulating gene expression to mitigate oxidative damage [35]. Thus, although *Pseudopithomyces* c174 is significantly enriched in NAFLD patients, its potential association with metabolites may act as a probiotic in the gut, providing a protective function. Phenylacetic acid, a novel microbial metabolite linked to aromatic amino acids metabolism, is significantly different between NAFLD patients and healthy controls in this study and relevant to liver diseases. Studies in primary human hepatocytes show that chronic exposure to phenylacetic acid triggers pathways leading to triglyceride accumulation [36]. Additionally, mice treated with this metabolite exhibit increased intrahepatic triglyceride content, suggesting a crucial role in the progression of NAFLD by promoting lipid accumulation in the liver [37]. Based on this, the fungi (*Aureobasidium* c170 and *Basipetospora* c193) that are positively correlated with phenylacetic acid may play a role in advancing NAFLD and therefore require careful attention. These results indicate that gut fungi play a significant role in the composition of human blood metabolites and the progression of NAFLD.

Fungi and bacteria form mixed polymicrobial communities in the gut, interacting through biofilms and exchanging nutrients, signaling molecules, and other biomolecules [38]. These interactions are not only fundamental to the microbiome’s structure and function but also have profound impacts o n the host’s metabolic health. Understanding the key fungal and bacterial taxa within this network and their interrelationships can reveal crucial insights into their roles in promoting health or contributing to metabolic diseases. In our study, despite minor differences in alpha and beta diversity within the gut mycobiome, significant variations in the co-abundance networks between bacteria and fungi indicate substantial changes in the interaction patterns of the gut microbiome associated with NAFLD. Existing research has demonstrated that Penicillium is one of the most representative fun gal genera in obese populations. It may be associated with obesity-related complications, such as NAFLD, and thereby as a key taxa in the gut of NAFLD patient [39]. Similarly, *Malassezia spp*., a fungus associated with inflammatory bowel disease (IBD), can produce metabolites that act as virulence factors to promote inflammation and exacerbate disease. This potential mechanism may also play a significant role in the development of NAFLD [40]. Mycotoxins from *Alternaria alternata* c42 may contribute to intestinal disorders by altering immune responses and disrupting the integrity of the epithelial barrier. These effects are critical in conditions such as inflammatory bowel disease and may also influence other metabolic disorders [41]. However, in the health control group, *Schizophyllum*-derived β-glucan beneficially modifies the gut environment, improving intestinal functions and offering protection against conditions such as constipation and obesity. These improvements are crucial in mitigating risk factors associated with metabolic disorders like NAFLD [42]. I n research on IBD, *Candida metapsilosis* has been identified as a beneficial gut fungus for treatment and prevention, effectively alleviating colitis symptoms in mice and potentially playing a crucial role in maintaining intestinal stability [43]. These findings indicate that although bacteria are more diverse and abundant in the gut microbiome, fungal species also play a significant role. Future efforts may need to focus on exploring the critical functions of the gut mycobiome within the intestinal environment.

Incorporating microbial taxa as sensitive biomarkers into machine-learning models for early disea se state prediction has shown immense potential [44, 45]. Previous studies have employed various indicators for predicting NAFLD, such as microbiome-metabolome combined models and bacteria l-clinical marker models, achieving AUC values ranging approximately from 0.58 to 0.80 [15, 46], indicating reasonably good effectiveness. However, the potential of gut fungi in predicting NAFL D remains an unexplored area in the research field. By integrating both bacterial and fungal specie s, using just 42 bacterial/fungal species, our model achieved an AUC of 0.772, demonstrating its superior predictive capability. However, within the entire predictive model, bacteria predominantly drive the predictions, with a greater number of bacterial taxa closely associated with NAFLD compared to fungi. Nonetheless, two fungal species also appear to play a role in the progression of NA FLD. Specifically, in the study mentioned above, *Alternaria alternata* c42 emerges as a key taxon in the NAFLD gut microbiome network, where it plays a significant role. Similarly, *Candida para psilosis*, a commensal fungus known for its extracellular lipase activity, is linked to diet-induced obesity. While it is primarily associated with conditions like gut fermentation syndrome, its metabolic byproducts, such as ethanol, could potentially contribute to liver inflammation and fat accumulation, thereby influencing the progression of NAFLD. However, our study has some limitations, including the absence of longitudinal data, which hinders our ability to monitor and analyze changes in the gut mycobiome of NAFLD patients over time. It is necessary to consider future work to address these issues. Additionally, the limited number of samples may have obscured differences between the microbial communities of the two groups. Therefore, more data are needed to validate the interactions between the metabolic system and the gut mycobiome in individuals with NAFLD.

## Conclusion

We conducted a metagenomic analysis of fecal samples from NAFLD patients to systematically describe the gut mycobiome of these individuals. Our study reveals significant alterations in the gut fungal community of NAFLD patients, which are associated with host immunity, inflammation, and toxin levels, ultimately impacting patient health. This research offers valuable insights for future mechanistic and clinical intervention studies. Additionally, due to the lack of detailed clinical information for these samples, we did not account for other potential risk factors for gut fungal infect ions, such as obesity and diabetes. Further datasets are needed to validate the influence of these factors.

## Acknowledgements

Not applicable.

## Author contributions

NZ, QLY, SHL, JY and CMJ contributed to the conception and design of the study. and DRW, G RX, YDG performed the data analysis and investigation. JL, JK and SSS drafted the manuscript. L C and SF revised the manuscript. All authors read the manuscript, contributed to the article, and approved the submitted version.

## Funding

Not applicable.

## Data availability

The raw metagenomic sequencing dataset for this study has been were downloaded from the NCB I Sequence Read Archive (SRA) under the accession ID PRJNA728908 and PRJNA686835 (only 4 control samples with ID SRR13279648, SRR13279753, SRR13279664, and SRR13279666).

## Declarations

## Ethics approval and consent to participate

Not applicable.

## Competing interests

All authors have no competing interests to disclose.

